# The disordered regions of SETD2 govern H3K36me3 deposition by regulating its proteasome-mediated decay

**DOI:** 10.1101/2020.06.05.137232

**Authors:** Saikat Bhattacharya, Ning Zhang, Hua Li, Jerry L. Workman

## Abstract

SETD2 is the sole methyltransferase that tri-methylates histone H3 at lysine 36 in mammals. It has an extended N-terminal region which is absent in its yeast homolog Set2. The function of this poorly characterized region in regulating SETD2 stability has been reported. However, how this region regulates SETD2 half-life and the consequences of the cellular accumulation of SETD2 is unclear. Here we show that the SETD2 N-terminal region contains disordered regions and is targeted for degradation by the proteasome. The marked increase in global H3K36me3 that occurs on the removal of the N-terminal segment results in a non-canonical distribution including reduced enrichment over gene bodies and exons. An increased SETD2 abundance leads to widespread changes in transcription and alternative splicing. Thus, the regulation of SETD2 levels through intrinsically disordered region-facilitated proteolysis is important to maintain the fidelity of transcription and splicing related processes.

## INTRODUCTION

The SET [**S**u(var)3-9, **E**nhancer-of-zeste and **T**rithorax] domain-containing enzyme, Set2/SETD2, is a crucial methyltransferase conserved from yeast to mammalian cells. It methylates lysine 36 of histone H3, a functionally important mark that suppresses cryptic transcription and is involved in DNA repair, pre-mRNA splicing and DNA methylation (Venkatesh et al. 2012)(Qin et al. 2017)(Kolasinska-zwierz et al. 2009)(Dhayalan et al. 2010)(Li et al. 2013)(Pfister et al. 2014). The H3K36me3 mark is deposited co-transcriptionally through the association of SETD2 with the large subunit of the RNA polymerase II, Rpb1, through its SRI (**S**et2-**R**pb1 **I**nteraction) domain. Besides histone H3 methylation, SETD2 also has non-histone substrates such as tubulin and STAT1 (Park et al. 2016)(Chen et al. 2017a). Consistent with its functionally important role, SETD2 deletion is embryonically lethal and mutations in SETD2 have been reported in many cancers, including CCRCC (Hu et al. 2010)(Zhu et al. 2014)(Duns et al. 2010)(Al Sarakbi et al. 2009)(Chiang et al. 2018)(Su et al. 2017)(Li et al. 2016). SETD2 depletion is clearly detrimental to cells.

SETD2 is functionally important, however, counterintuitively the SETD2 protein doesn’t accumulate in human cells. We and others have shown that SETD2 is robustly degraded by the **u**biquitin-**p**roteasome **s**ystem (UPS)(Bhattacharya and Workman 2020)(Zhu et al. 2016). Notably, SETD2 has an N-terminal region that is absent in its yeast homolog Set2. We previously reported that this unique region when removed, leads to the stabilization of SETD2 and a marked increase in the level of H3K36me3 mark which happens uncharacteristically in an RNA Poll II independent manner (Bhattacharya and Workman 2020). However, how this N-terminal segment regulates SETD2 stability is unclear. Also, the possible consequences of the cellular accumulation of SETD2 have not been investigated.

The correct turnover of proteins is important for maintaining cellular homeostasis and function. The UPS is the major proteolytic proteasomal pathway in eukaryotic cells that mediates the controlled and selective degradation of proteins (Cooper 2000). Protein substrates are targeted for degradation through their polyubiquitination by ubiquitin ligases (Kommander and Rape 2012)(Ravid and Hochstrasser 2008). Besides ubiquitination, the role of intrinsic structural features of substrates in influencing their half-life is emerging. One such feature is the presence of **i**ntrinsically **d**isordered **r**egions (IDRs) which are segments in a protein that do not adopt a defined 3D structure (van der Lee et al. 2014). IDRs can be important in regulating protein turn over and are known to regulate the half-life of proteins such as ODC, Rpn4, TS, p53, p21, c-Jun and α-synuclein (Gödderz et al. 2011)(Melo et al. 2011)(Yoon et al. 2012)(Erales and Coffino 2014). IDRs are often involved in diseases such as cancer, cardiovascular disease, and neurodegenerative diseases (Uversky et al. 2008).

Here we show that the SETD2 protein has intrinsically disordered regions that regulate its half-life through proteasomal degradation. In the absence of SETD2 degradation, spurious deposition of H3K36me3 occurs in a non-canonical manner that is accompanied by widespread transcription as well as alternative splicing aberrations.

## RESULTS

### Deletion of the N-terminal region of SETD2 leads to changes in transcription and splicing

SETD2 is a large protein with 2564 amino acids. The C-terminal region of the protein (1404-2564 residues) harbor all the characterized domains that are conserved with ySet2. However, SETD2 has an N-terminal segment (1-1403 residues) not found in ySet2 [Figure 1a]. We have previously shown that the removal of the N-terminal region of SETD2 leads to its stabilization and results in a marked increase in H3K36me3 levels (Bhattacharya and Workman 2020). Next, we wanted to investigate whether the removal of the SETD2 N-terminal region leads to transcriptome changes.

**Figure 1:**
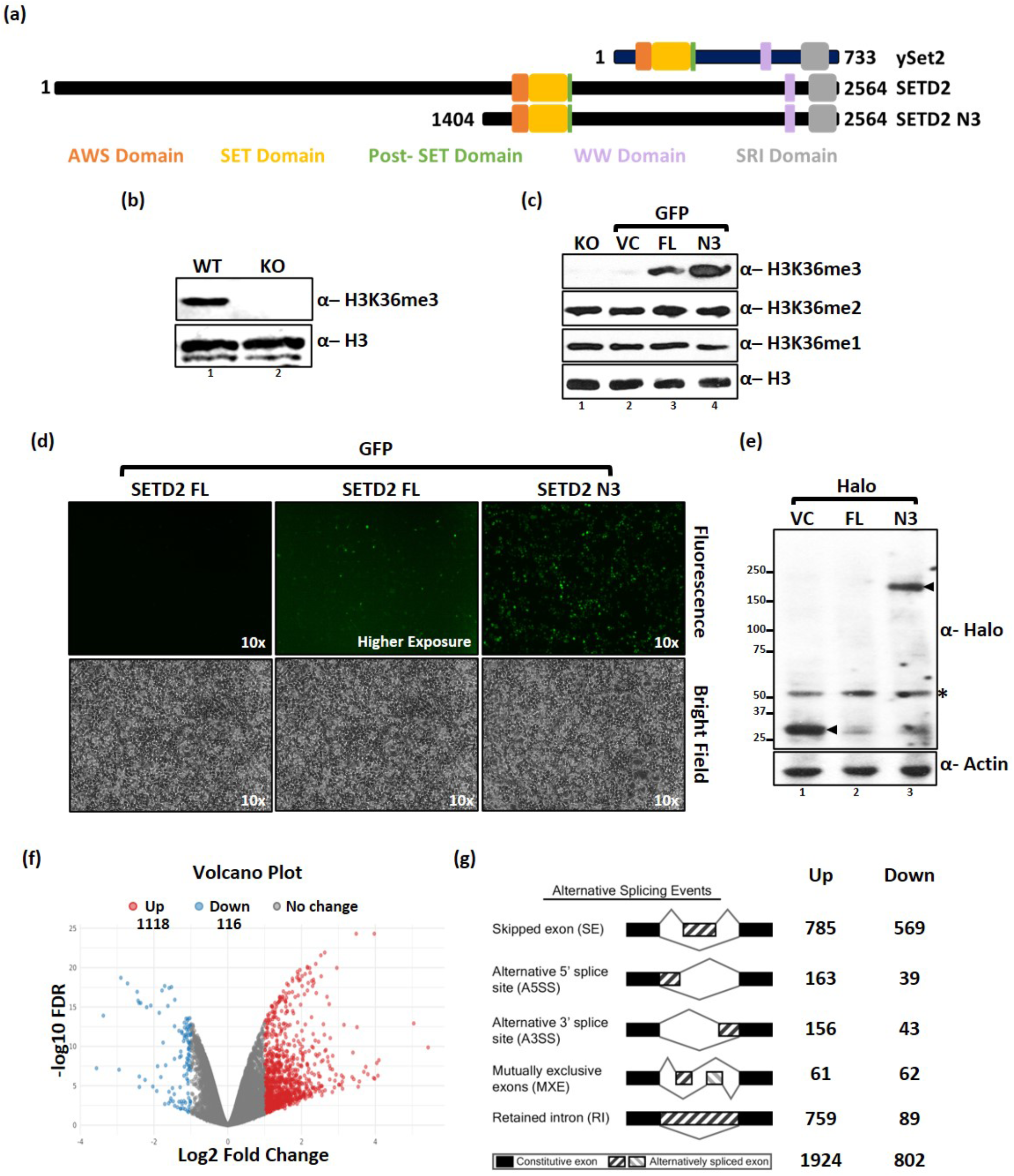
Removal of the SETD2 N-terminus leads to its increased abundance and changes in the transcriptome. (a) Cartoon illustrating the length and domains of ySet2/SETD2. (b, c, e) Western blot of whole-cell lysates probed with the depicted antibodies. setd2Δ 293T (KO) cells were transfected with Halo-vector control (VC), SETD2 full-length (FL), or SETD2 N3 (N3) and lysates were prepared 72 hrs post-transfection. * denotes non-specific band. The expected bands for the target are depicted by arrows. (d) Microscopy images showing the expression of GFP-SETD2 constructs in 293T cells. (f) Volcano plot showing the expression changes occurring on SETD2 N3 expression as compared to SETD2 FL. Each dot represents a gene. RNA was isolated 72 hours post-transfection. (g) RMATS plot depicting the differential AS events in SETD2 N3 rescued KO cells as compared to SETD2 FL rescued ones. The cartoon was taken from http://rnaseq-mats.sourceforge.net/.

To test this, constructs were made to express GFP- or Halo-tagged full-length (SETD2 FL) and the C-terminal segment of SETD2 (1404-2564, SETD2 N3) under the control of a CMV promoter. Recombinant expression of SETD2 has been previously used to investigate the function of the protein (Zhang et al. 2020)(Chiang et al. 2018)(Zhu et al. 2016)(Carvalho et al. 2014)(Chen et al. 2017a). The constructs were introduced in *setd2Δ* 293T (KO) cells in both the alleles of the endogenous SETD2 gene were disrupted (Hacker et al. 2016). Consistent with the role of SETD2 as the sole H3K36me3 depositor in humans, in the KO cells the H3K36me3 mark was not detected in the whole-cell lysates by immunoblotting [Figure 1b]. 72 hours post-transfection with the GFP tagged constructs, whole-cell lysates were prepared and analyzed by western blotting. Consistent with our previous report, the expression of SETD2 N3 in KO cells led to a marked increase in the H3K36me3 level as compared to the rescue with SETD2 FL [Figure 1c] (Bhattacharya and Workman 2020). The other two H3K36 methyl marks, H3K36me1 and H3K36me2, largely remained unchanged [Figure 1c]. The expression of the empty vector control (VC) did not alter H3K36 methylation as expected [Figure 1c]. Further, microscopy of cells expressing GFP-tagged constructs and western blotting of lysates of cells expressing Halo-tagged proteins revealed that the expression of SETD2 N3 is markedly higher than that of SETD2 FL like we have previously reported [Figure 1d, e] (Bhattacharya and Workman 2020).

Subsequently, we used these cells to perform gene expression analysis. Expression of SETD2 FL and SETD2 N3 led to pronounced transcriptome changes in the KO cells [supplementary information S1]. Strikingly, a comparison of SETD2 N3 versus SETD2 FL-expressing cells revealed that a total of 1234 genes exhibited significant differential expression [Fold change (FC) ≥2, false discovery rate (FDR) ≤0.05] [Figure 1f]. The majority of these genes (1118 genes) were upregulated on SETD2 N3 expression and only 116 were downregulated in comparison to SETD2 FL. Noticeably, Gene Ontology Enrichment Analysis revealed that the upregulated genes are not enriched in any pathway, suggesting, that the removal of the N-terminal region of SETD2 leads to widespread changes in transcription without affecting any specific pathway. Analysis of our transcriptome data to look for possible differences in splicing also revealed widespread changes in cells expressing SETD2 N3 versus SETD2 FL. A total of 2726 differential alternative splicing (AS) events (FDR ≤0.05) were identified. Out of these, 1924 events were upregulated and 802 were downregulated upon SETD2 N3 expression [Figure 1g]. The changes belonged to the different kinds of splicing events, such as skipped exon, alternative 5’ splice site, alternative 3’ splice site, mutually exclusive exons, and retained intron. Most changes belonged to skipped exon and retained intron events. Also, the transcript levels of SETD2 FL and SETD2 N3 were not significantly different in our RNA-seq data, suggesting that the differences observed are not due to the differences in the transcription of the two constructs.

Collectively, our data suggest that removal of the N-terminal region of SETD2 leads to widespread changes in the transcriptome of mammalian cells.

### SETD2 has multiple long intrinsically disordered regions

We previously showed that SETD2 is degraded by the ubiquitin-proteasome system (UPS)(Bhattacharya and Workman 2020). We next wanted to learn more about the UPS-mediated decay of SETD2. We checked the contribution of the E3 ubiquitin ligase SPOP, the anaphase-promoting complex (APC) and the PEST motif on SETD2 degradation (Zhu et al. 2016)(Dronamraju et al. 2018)(Rechsteiner and Rogers 1996) [please see supplementary information S2 and S3 and the associated results]. Our results agreed with the previously published data that the SPOP in part regulates SETD2 stability, but also indicated that the post-translational control of SETD2 abundance is more complicated than previously thought and may contain several redundancies.

We previously reported the N-terminal region of SETD2 regulates its stability (Bhattacharya and Workman 2020). Notably, the NCBI Conserved Domain Search (Marchler-Bauer and Bryant 2004) revealed that the SETD2 N-terminus lacks any known domains or motifs and did not yield any clue about its possible function. Also, the SETD2 N-terminus has very little sequence similarity with other known protein sequences. Consequently, homology modeling of the structure using SWISS-MODEL (Waterhouse et al. 2018) failed. Strikingly, the sequence is also 81% disordered based on Robetta prediction (http://robetta.bakerlab.org/). This was of interest because long N-terminal and internal disordered segments contribute to short protein half-life *in vivo* (Tompa et al. 2008)(van der Lee et al. 2014). This prompted us to perform a deeper analysis of the disordered regions in SETD2 protein.

Prediction of the disordered region in SETD2 was performed using IUPRED2 (Mészáros et al. 2018). IUPred provides a score between 0 and 1 for every residue, with a score of greater than 0.5 means that it is more likely that the residue is a part of a disordered region. Strikingly, 1546 of the 2564 (60.29%) residues of the SETD2 protein returned a score of >0.5 as compared to a Halo control (0%) [Figure 2a, b, c, f]. Furthermore, 67.6% of residues in the N-terminus scored >0.5 as compared to 51.4% of residues of C-terminus. Next, we looked for long disordered regions (>30 residues) in SETD2. The length was based on findings that a critical minimum length of ~30 residues allows a disordered terminus of a ubiquitinated substrate to efficiently initiate proteasomal degradation (Da Fonseca et al. 2012)(Lasker et al. 2012b) (van der Lee et al. 2014). Strikingly, 17 such stretches were found in SETD2 out of which 11 were in the N-terminal region, 5 in C-terminal fragment, and 1 overlapping between and the N and C-terminal segments [Figure 2f].

**Figure 2:**
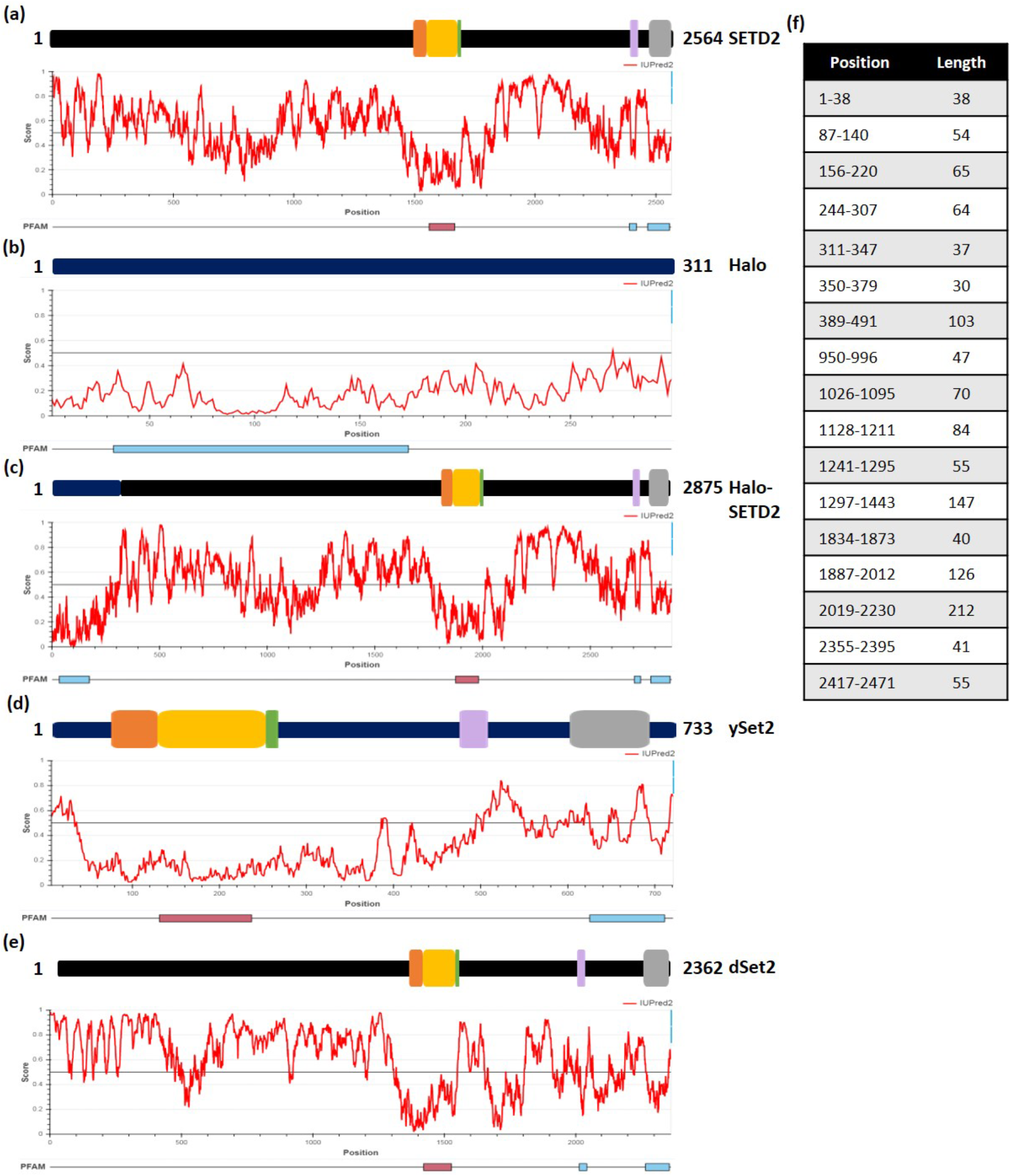
SETD2 has intrinsically disordered regions. Various protein sequences were subjected to disordered region prediction using IUPRED2A. See the text for more details.

Our findings regarding stability and disordered region prediction in SETD2 suggest the possibility that the disordered regions of SETD2 may govern its half-life.

### Multiple disordered segments of SETD2 have a combined effect on its half-life

Proteins with several internal disordered segments have shorter half-lives than proteins with only one such segment (van der Lee et al. 2014). SETD2 undergoes robust degradation and is predicted to have numerous disordered segments throughout. We speculated that if the multiple disordered segments of SETD2 collectively enhance its proteolysis, then the expression of the shorter fragments will be higher than the longer ones. To test this, a series of constructs were made to express Halo-tagged N- or C-terminal truncations of SETD2 in 293T cells [Figure 3a, b]. The use of recombinant truncation mutants to determine the region of protease sensitivity in a protein is a commonly used approach and has been successfully used (Sano et al. 2007)(Zhu et al. 2016). Western blotting of whole-cell extracts with an anti-Halo revealed that for both the N and C-terminal truncations, SETD2 protein expression anti-correlated with its length [Figure 3c, d]. Also, in our previous report we showed that three non-overlapping fragments of Halo-SETD2; N1a (C4), N1b (504-1403) and C (N3) expressed robustly in 293T cells, indicating that smaller fragments of SETD2 accumulate more than SETD2 FL, consistent with our hypothesis (Bhattacharya and Workman 2020).

**Figure 3:**
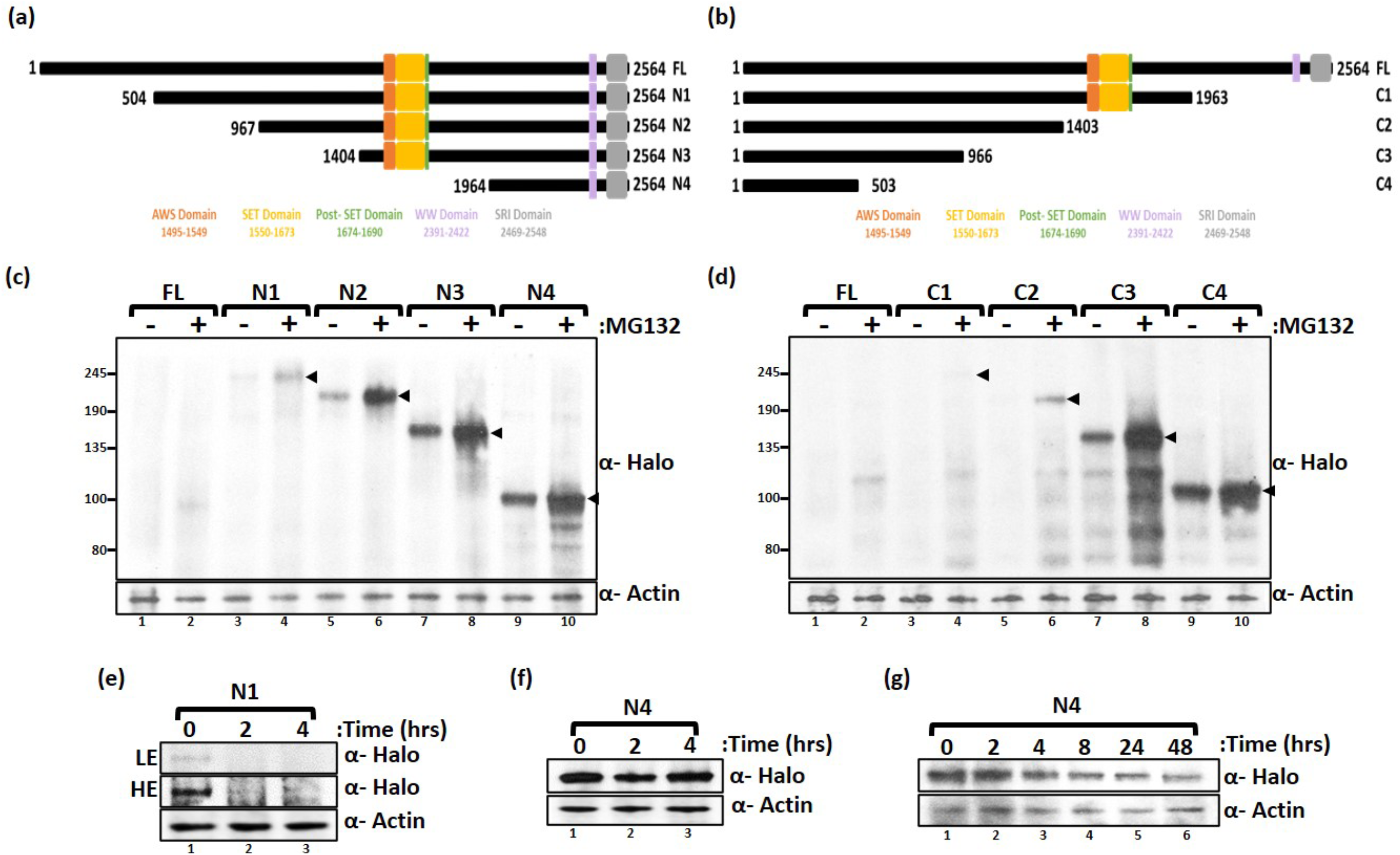
Deleting SETD2 fragments leads to its stabilization. (a, b) Cartoon illustrating the N and C-terminal truncations of SETD2 along with the known domains. (c, d) Western blot of whole-cell lysates probed with the depicted antibodies. Lysates of 293T cells expressing Halo-SETD2 constructs were prepared after 12 hrs of MG132 (10 μM) treatment. The expected band for the target protein is depicted by arrows. (e, f, g) Western blot of whole-cell lysates probed with the depicted antibodies. Lysates of 293T cells expressing Halo-SETD2 constructs were prepared after cycloheximide (10 μM) treatment. The duration of the treatment is shown in the figure. HE-High Exposure, LE-Low Exposure.

To confirm that the differences observed in the expression are indeed due to the shorter half-life of larger SETD2 fragments, a time-chase experiment was performed to monitor the expression of Halo-N1 and N4 upon the treatment of cells with cycloheximide (CHX), a protein translation inhibitor. Post 2 hours CHX treatment, Halo-N1 could no longer be detected in the whole-cell lysates whereas no appreciable decrease was observed for Halo-N4 even after 4 hours [Figure 3e, f]. Notably, Halo-N4 could be readily detected even after 48 hours of CHX treatment [Figure 3g] demonstrating that the shorter fragment N4 has a longer half-life in comparison to N1. Importantly, the observed differences in the half-life of the proteins along with our RT-PCR and RNA-seq data validate that the dissimilarities in the expression of the SETD2 constructs are indeed due to the differences in protein stability and is not due to experimental variations like transfection efficiency.

From these experiments, no specific region emerged in the SETD2 protein that is particularly targeted for UPS-mediated decay and confirmed that the turnover of SETD2 is regulated by factors besides those that were previously reported. Furthermore, consistent with the report that IDRs have a combined effect on protein half-life, multiple regions of SETD2 were found to co-operatively regulate its proteolysis.

### The IDR-rich N-terminus of SETD2 can reduce the half-life of ySet2

We previously showed that the ySet2 localized to the nucleus of mammalian cells and its expression was significantly higher than SETD2 FL and is comparable to SETD2 C (N3) with which it shares conservation (Bhattacharya and Workman 2020). Disordered region prediction revealed that overall ySet2 is a well-ordered protein with a much lower proportion of residues (24.69%) predicted to be disordered as compared to its homolog SETD2 [Figure 2d]. We speculated that the disparity in expression between ySet2 and SETD2 could be due to the differences in the disordered region abundance between the proteins. We next wanted to test whether the N-terminal disordered segment of SETD2 can destabilize the ySet2 protein when fused to it [Figure 4a].

**Figure 4:**
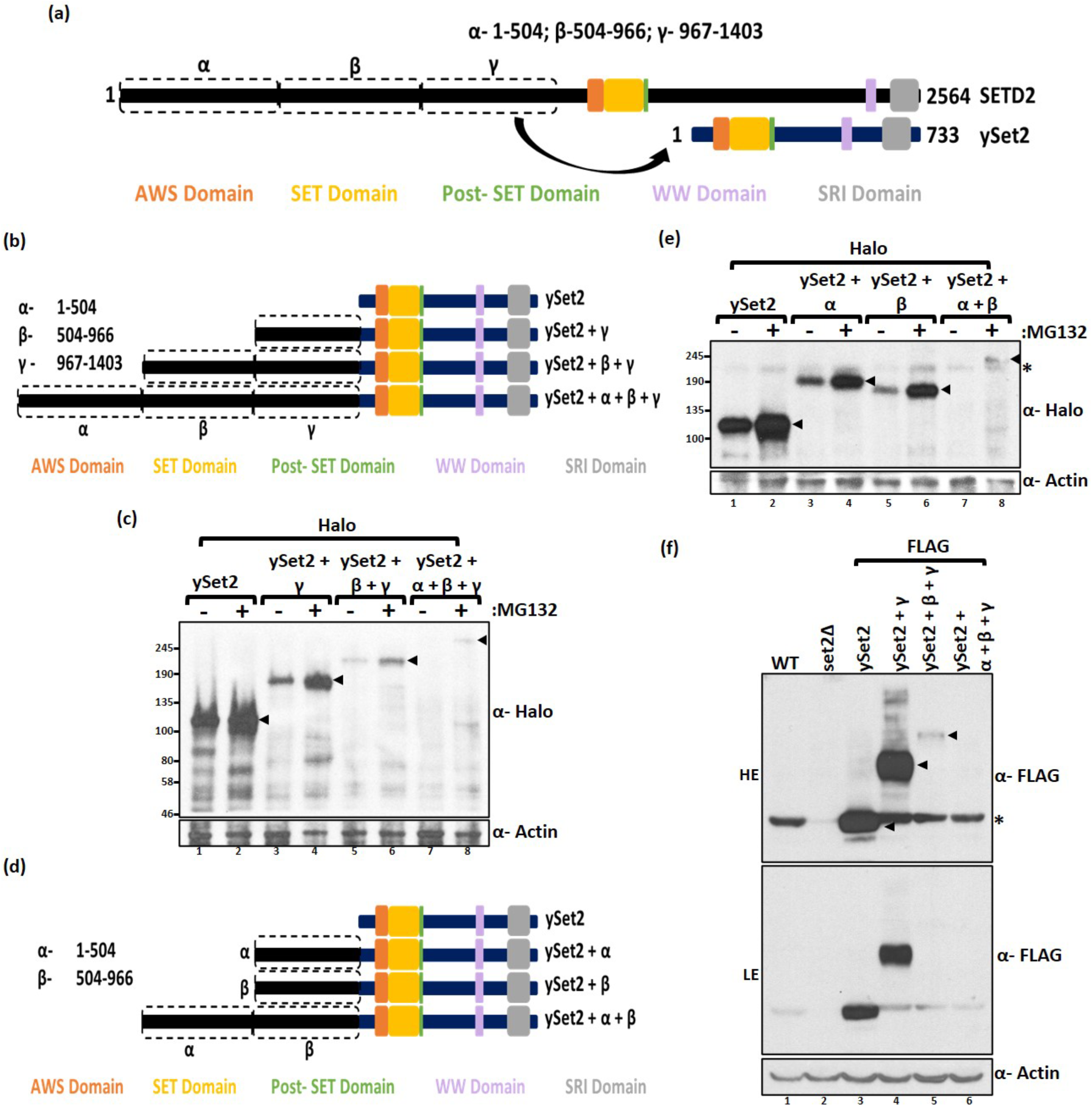
The SETD2 N-terminus can destabilize ySet2. (a, b, d) Cartoon illustrating the chimeric ySet2 constructs made by fusing with the segments of N-terminus of SETD2. (c, e) Western blot of whole-cell lysates probed with the depicted antibodies. Lysates of 293T cells expressing Halo-ySet2 constructs were prepared after 12 hrs of MG132 (10 μM) treatment. The expected band for the target protein is depicted by arrows. (f) Western blot of whole-cell lysates of yeast BY4741 expressing FLAG-ySet2 constructs (using pRS416) probed with the depicted antibodies. HE-High Exposure, LE-Low Exposure, *-non-specific.

For this, constructs were made to express Halo-tagged chimeric proteins that have increasing stretches of the N-terminal segment of SETD2 (α: 967-1403, β: 504-1403, and γ: 1–1403) fused upstream of ySet2 [Figure 4b]. The expression of these constructs was tested in 293T cells by probing whole-cell lysates with an anti-Halo antibody. A progressive decrease in the accumulation of chimeric ySet2 was observed with the increasing length of the N-terminal segment and all the chimeras displayed sensitivity to MG132 treatment [Figure 4c]. Next, we wanted to test whether the destabilization brought about by the SETD2 N-terminus is influenced by the order in which the fragments are added upstream of ySet2. For this, the expression of a different set of chimeric constructs was tested in 293T cells [Figure 4d, e]. The addition of α and β destabilized ySet2 and fusing α + β destabilized it further, even in the absence of fragment γ. The data also suggests that the SETD2 fragments co-operatively regulate stability which is consistent with the idea that IDRs have a combined effect on protein half-life.

The architecture of the proteasome is conserved from yeast to mammals (Da Fonseca et al. 2012)(Lasker et al. 2012a). Consistent with that, we previously showed that ySet2 responds to MG132 treatment in 293T cells, suggesting, that it is targeted for degradation through UPS in human cells too (Bhattacharya and Workman 2020). We wondered whether the IDR-rich SETD2 N-terminal segment from humans can enhance ySet2 proteolysis in yeast. To test this, the expression of FLAG-tagged WT and chimeric ySet2 proteins was checked in the yeast strain BY4741. The expression was scored by probing the whole-cell extracts with an anti-FLAG antibody. Although the addition of the SETD2 fragment γ (967-1403) did not have a large impact, the addition of β and α + β had a drastic destabilization effect on the ySet2 protein [Figure 4f].

Collectively, our results demonstrate that the IDR rich N-terminus of the SETD2 protein has a destabilization effect on both SETD2 and its yeast homolog ySet2. Importantly, they also suggest evolutionary conservation of the role of the N-terminal region of SETD2 in bringing about the degradation of a protein without the need for specific E3 ubiquitin ligases. This is key evidence for IDR-mediated degradation as IDRs directly act as efficient sites for initiating proteasomal degradation (van der Lee et al. 2014).

### High levels of SETD2 leads to a non-canonical H3K36me3 distribution

The removal of the N-terminal region leads to a marked increase in the H3K36me3 level. We wondered whether the increased SETD2 accumulation also leads to an altered distribution of the H3K36me3 mark. For this, first, spike-in normalized H3K36me3 ChIP-Seq of WT and KO cells was performed. Metagene analysis revealed a clear enrichment of H3K36me3 within the coding region of the genes in the WT cells [supplementary information S4a]. As expected, a similar pattern was not observed in KO cells that lack SETD2 and hence, H3K36me3 [supplementary information S4a]. Additionally, a metagene analysis of H3 normalized H3K36me3 of protein-coding genes highlighted that H3K36me3 is greatly enriched on the highly expressed genes as compared to the lowly expressed ones, consistent with the idea that the mark is associated with transcriptionally active genes [supplementary information S4b]. Furthermore, a closer inspection of the H3 normalized H3K36me3 distribution within the coding region revealed that H3K36me3 is more enriched over exons than introns [supplementary information S4c].

Next, a ChIP-Seq of H3K36me3 was performed post introduction of GFP-SETD2 constructs in KO cells and their distribution was compared. On rescue of KO cells with the SETD2 constructs, the H3K36me3 mark continued to be enriched within the gene bodies [Figure 5a]. Notably, this was true even for SETD2 N3ΔSRI and is consistent with the reports that association with RNA Pol II is not required for Set2 association with highly transcribed genes (discussed in the Discussion section). Also, the H3K36me3 level positively correlated with increased gene expression [supplementary information S4d]. However, a closer inspection of the distribution reveals that it is skewed towards the 5’ end of the genes in cells rescued with N3 as compared to FL [Figure 5a]. Furthermore, a global analysis of the H3K36me3 peaks revealed that there is a higher deposition of H3K36me3 at the intergenic regions post-rescue with SETD2 N3 as compared to FL [Figure 5b]. In addition, there was an aberrant deposition of H3K36me3 on genes such as CLIC4 although no change in its expression was observed in our RNA-seq data [Figure 5c]. Furthermore, the 5’ skewing observed in the metagene analysis was exemplified in genes such as ARID1A [Figure 5d]. Strikingly, a loss of H3K36me3 enrichment over exons was observed in the ARID1A gene in SETD2 N3 and N3ΔSRI expressing cells as compared to SETD2 FL expressing ones. Indeed, a global analysis revealed that the ratio of average exon signal divided by the average intron signal was lower in N3 rescued cells than FL rescued cells [Figure 5e]. This suggests that on an increased accumulation of SETD2, the enrichment of H3K36me3 over exons decreases. Next, we analyzed the H3K36me3 exon enrichment analysis of the genes showing differential AS events described in Figure 1g. Interestingly, an anticorrelation of exon enrichment score was observed with splicing event upon N3 expression, meaning where splicing was down regulated a higher enrichment score was seen [Figure 5f]. However, this negative correlation was not seen in FL expressing cells in which a constant exon enrichment value of 1.5 was observed over the same genes [Figure 5f]. Also, the H3K36me3 levels were higher on the genes that showed increased splicing in N3 versus FL [Figure 5g].

**Figure 5:**
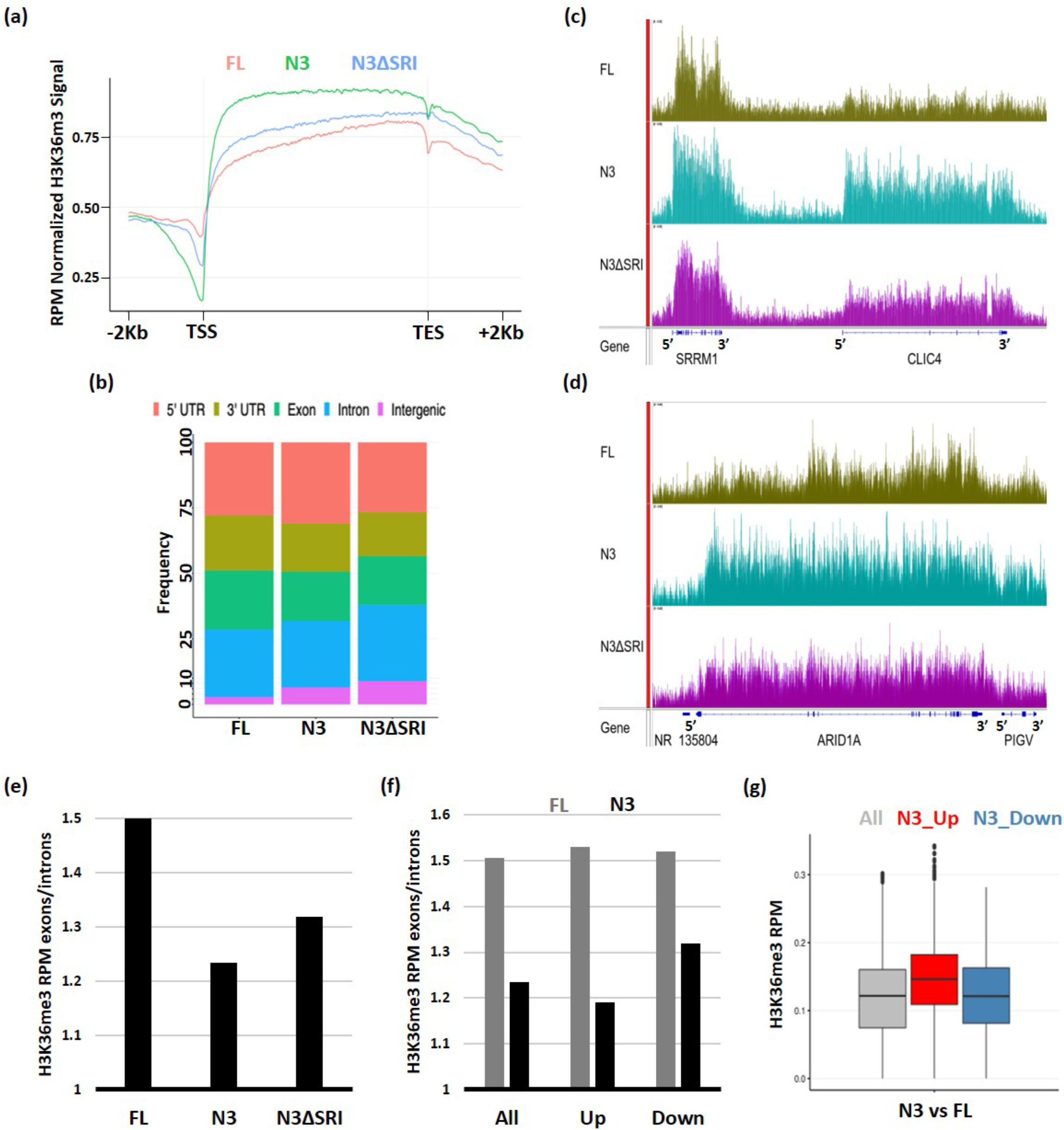
Increased SETD2 abundance leads to a non-canonical H3K36me3 deposition. (a, b) Metagene plot and histogram depicting the distribution of H3K36me3. (c, d) Genome browser view depicting the peaks of H3K36me3. The blue solid blocks on the X-axis denotes the exons. (e) Bar graph showing the H3K36me3 enrichment over exons as compared to introns. (f) Bar graph showing the H3K36me3 enrichment over exons as compared to introns of genes that show an increase (Up) or decrease (Down) in splicing events in N3 as compared to FL. “All” is the overall enrichment of H3K36me3 over all genes. (g) Box plot showing the H3K36me3 of genes that show differential alternative splicing upon SETD2 N3 expression as compared to SETD2 FL.

Thus, in the absence of the N-terminal rich IDRs, there is a reduced initiation of UPS-mediated decay of SETD2 resulting in its high abundance, reduced Pol II dependency, and leading to global changes in the canonical distribution pattern of H3K36me3 [Figure 6].

**Figure 6:**
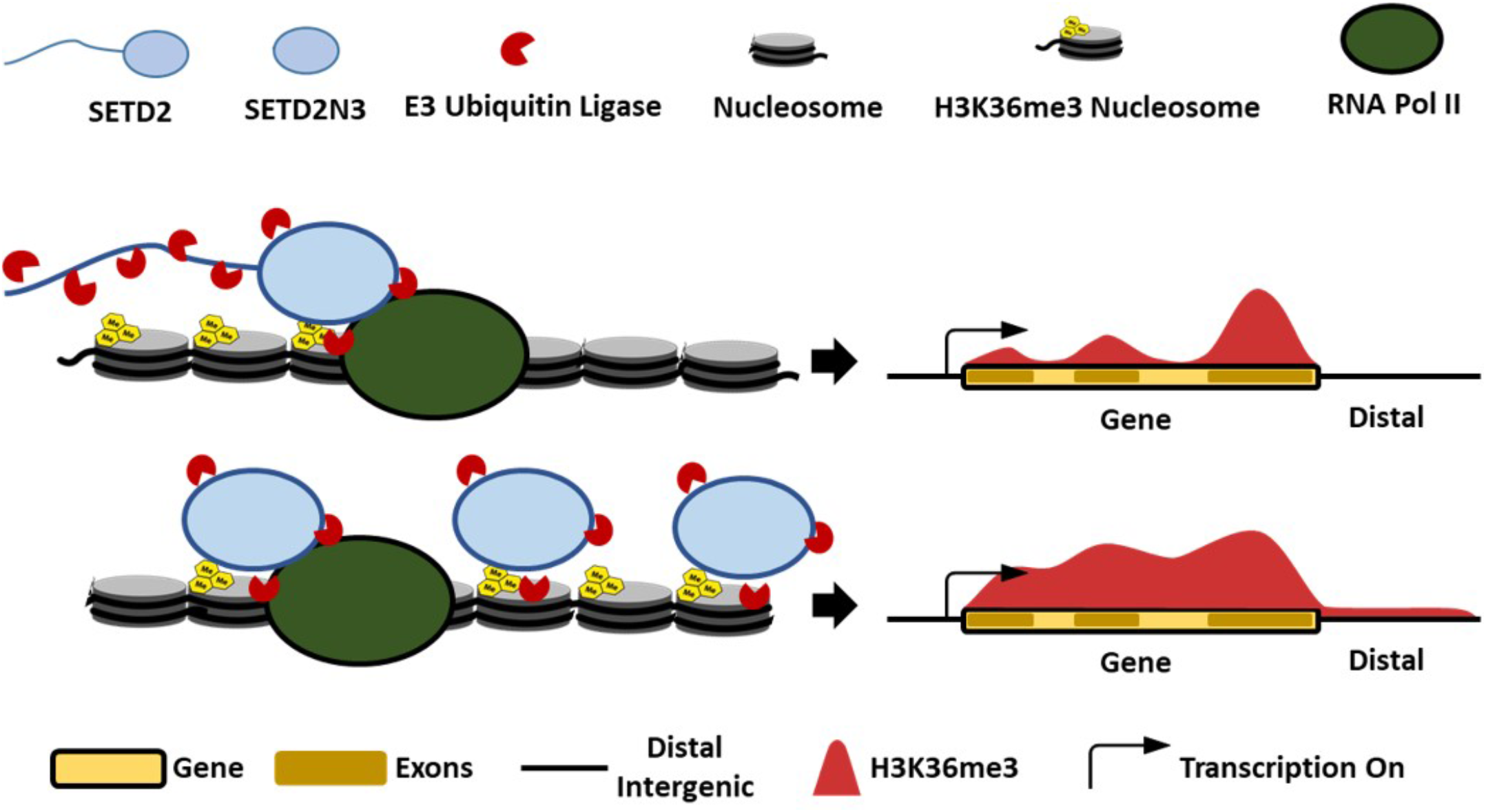
Removal of the N-terminal region from SETD2 stabilizes the protein and results in a non-canonical H3K36me3 distribution.

## DISCUSSION

Although we and others have reported the role of the proteasome in regulating SETD2 stability, how SETD2 degradation occurs so robustly was not clear. Furthermore, it was not clear why the cellular half-life of SETD2 is so tightly regulated. We not only show that multiple regions of SETD2 are targeted for proteasomal degradation, but also found that the disordered segments of SETD2 are important in this process. Importantly, our work clearly illustrates the importance of the N-terminal segment of SETD2 in governing its appropriate function.

### SETD2 IDRs possibly act as efficient sites for initiating proteasomal degradation

An analysis of the functional annotations of proteins with long disordered segments revealed enrichment for associations with regulatory and transcription functions (van der Lee et al. 2014). This suggests that the IDR-mediated regulation of protein half-life that we discovered for SETD2 might be a prevalent mechanism employed by cells to govern essential processes. Disordered segments lack motifs and act directly to regulate protein half-lives by forming initiation sites for degradation by the proteasome (van der Lee et al. 2014)(Zhao et al. 2010)(Prakash et al. 2004)(Verhoef et al. 2009). The access to the proteasome is an important step that regulates the half-life of substrates (Goldberg 2003)(Hershko and Ciechanover 1998). The catalytic residues are situated deep within the proteasome core particle and are only accessible through a long narrow channel. A terminal disordered segment of 30 residues or an internal disordered segment of at least 40 residues can span twice this distance and thus, could be cleaved by the core particle (van der Lee et al. 2014). Thus, proteins such as SETD2 that have 17 long disordered segments are expected to be processed quickly due to the efficient initiation of degradation. In addition to that, the disordered segments are spread across the length of the protein and co-operatively leads to the robust degradation of SETD2.

### Variations in the N-terminal region might tune SETD2 half-life

The gain or loss of disordered segments may be an important contributor to the degradation rate of proteins during evolution (van der Lee et al. 2014). Variation in IDRs might be a mechanism for the divergence of half-life among orthologous proteins. IDRs are not required to attain a specified three-dimensional conformation to exert their functional effect and hence, can undergo mutations without greatly affecting their functionality. Such variation in disordered segments may provide an evolutionary mechanism for fine-tuning protein turnover rates. These forces might lead to inter- as well as intra-species divergence of protein half-life. Analysis of Drosophila SETD2 revealed large and abundant disordered regions [Figure 2e]. The differences in the degree of disorder between Drosophila and human SETD2 proteins might lead to differences in their half-lives much like what we observed between SETD2 and ySet2. Differences in half-lives between Set2 homologs might be required to adjust for the differences in the deposition mechanism of H3K36 methylation that we speculated previously (Bhattacharya and Workman 2020).

In addition to inter-species differences, intra-species variation in the length of IDRs may arise through mechanisms such as repeat expansion, AS, and alternative transcription start sites. Interestingly, in the ENSEMBL database, SETD2 has 9 splice variants out of which three are protein-coding. One of those with transcript ID ENST00000638947.1 codes for only a 591 residue segment of SETD2 that, importantly, contains the catalytic SET domain. This segment is trimmed off the IDRs present in full-length SETD2 and might be expected to accumulate to a higher level with inadvertent consequences as discussed below. Furthermore, differences in protein turnover may disturb protein abundance and could lead to disease (Ciechanover 2007)(Yang et al. 2012). Strikingly, many missense and truncation mutations that can potentially alter half-life, are found in SETD2 in cancers [supplementary information S5].

### SETD2 over-abundance might have inadvertent consequences

Besides the challenge of dealing with undesired protein aggregation that we previously discussed (Bhattacharya and Workman 2020), overexpression of SETD2 leads to widespread changes in the transcriptome which is consistent with the observed global increase and the altered distribution of H3K36me3. The list includes genes like NEAT1, MALAT1, MMP10, MMP12, and MMP13 that not only are functionally important but also has disease relevance (West et al. 2014)(Hirata et al. 2015)(Chen et al. 2017b)(Vaalamo et al. 1998). Furthermore, genes like the H3K9 demethylase KDM3A (FC 2.005, FDR 8.28e-14), the splicing factor U2AF1 (FC 2.009, FDR 1.6924e-5), 40 genes from the ZNF family were upregulated amongst others that have well established functional importance.

Besides affecting the transcriptome, the aberrant H3K36me3 can result in the mistargeting of important epigenetic regulators like DNMT3a, MutSα, and MORF (Bhattacharya and Workman 2020). The expression of SETD2 shorter splice variant with transcript ID ENST00000638947.1 might lead to similar perturbations in the H3K36me3 profile to what was observed upon expression of SETD2 N3. Also, the missense and truncation mutations found in SETD2 in cancers can potentially alter half-life, leading to aberrant H3K36me3 deposition [supplementary information S5].

### Pol II association is required for enhancing SETD2 activity but not for its recruitment

We previously showed that when the expression level of SETD2 is high, it has reduced dependency on RNA Pol II for H3K36me3 deposition (Bhattacharya and Workman 2020). Strikingly, the ChiP-Seq data of H3K36me3 suggests that RNA Pol II association is not required for chromatin recruitment of SETD2 as previously assumed, but rather the interaction is required for the activation of SETD2 enzymatic activity like we previously speculated (Bhattacharya and Workman 2020). Studies in yeast have also shown that the Pol II association is required for Set2 enzymatic activation (Wang et al. 2015). Notably, the Set2ΔSRI ChiP profile shows an enrichment on the coding sequence of genes that is skewed towards the 5’ end much like what we found for H3K36me3 when SETD2 N3ΔSRI is introduced in *setd2Δ* cells (Suzuki et al. 2016). Possibly, Set2/SETD2 can continue to engage with the transcription elongation complex even in the absence of its interaction with RNA Pol II. In future studies, it will be interesting to determine how the gene body enrichment of H3K36me3 and Set2/SETD2 occurs even without the Pol II association.

## MATERIALS AND METHODS

### Plasmids

SETD2-HaloTag^®^ human ORF in pFN21A was procured from Promega. Deletion mutants of SETD2 were constructed by PCR (Phusion polymerase, NEB) using full-length SETD2 as a template and individual fragments were cloned. All constructs generated were confirmed by sequencing. pCDNA3-ySet2 was procured from Addgene.

### Cell line maintenance and drug treatment

HEK293T cells were procured from ATCC. Cells were maintained in DMEM supplemented with 10% FBS and 2 mM L-glutamine at 37 °C with 5% CO_2_. MG132 (Sigma) was added at a final concentration of 10 μM for 12 hours. Cycloheximide (Sigma) was added at a final concentration of 10 μM. Transfections were performed at cell confluency of 40% using Fugene HD (Promega) using a ratio of 1:4 of the plasmid (μg) to transfection reagent (μl).

### Isolation of total RNA and PCR

Total RNA was extracted from cells as per the manufacturer’s (Qiagen) instructions. It was further treated with DNaseI (NEB) for 30 min at 72 °C to degrade any possible DNA contamination. RNA (2 μg) was subjected to reverse transcription using the QScript cDNA synthesis mix according to the manufacturer’s instructions. cDNAs were then amplified with the corresponding gene-specific primer sets. For RTPCR, PCR was conducted for 24 cycles using the condition of 30 s at 94 °C, 30 s at 60 °C and 30 s at 72 °C. The PCR products were analyzed on a 1% agarose gels containing 0.5 μg/ml ethidium bromide. The sequence of oligos is in supplementary information 6.

### Histone isolation and immunoblot analysis

Histones were isolated and analyzed as described previously (Bhattacharya et al. 2017). For immunoblotting, histones were resolved on 15% SDS– polyacrylamide gel, transferred to PVDF membrane and probed with antibodies. Signals were detected by using the ECL plus detection kit (ThermoFisher).

### Antibodies

H3K36me3 (CST, 4909S), H3 (CST, 9715S), Halo (Promega, G9211), β-actin (Abcam, ab8224), SETD2 (Abclonal, A3194), FLAG (Sigma-Aldrich, A8592).

### ChIP

Cells were cross-linked by 1% formaldehyde for 10 mins, and then quenched in 125 mM glycine for 5 mins. After washing with cold 1x PBS thrice, cells were harvested by scraping and pelleted down by centrifugation. The cell pellet was resuspended in swelling buffer (25 mM HEPES pH 8, 1.5 mM MgCl_2_, 10 mM KCl, 0.1% NP40, 1 mM DTT, protease inhibitor cocktail), kept in ice for 10 mins and then dounced. The nuclear pellet was obtained by centrifugation and resuspended in sonication buffer (50 mM HEPES pH 8, 140 mM NaCl, 1 mM EDTA, 1% Triton X 100, 0.1% Na-deoxycholate, 0.1% SDS, protease inhibitor cocktail), followed by sonication on ice for 12 cycles (30% amplitude, 10 secs on / 60 secs off) using a Branson Sonicator. For spike-in normalization, the spike-in chromatin and antibody were added in the reaction as per the manufacturer’s recommendation (Active Motif). The chromatin was incubated with antibodies at 4 °C overnight and then added to 30 μl of protein G-Dyna beads (Thermo Fisher Scientific) for an additional 2 hours with constant rotation. The beads were extensively washed, and bound DNA was eluted with elution buffer (50 mM Tris-HCl pH 8, 5 mM EDTA, 50 mM NaCl, 1% SDS) and reverse-crosslinked at 65 °C overnight. DNAs were purified using the QIAquick PCR purification kit (Qiagen) after the treatment of proteinase K and RNase A.

### High throughput sequencing

Sequencing libraries were prepared using High Throughput Library Prep Kit (KAPA Biosystems) following the manufacturer’s instructions. The library was sequenced on an Illumina HiSeq platform with paired reads of 75 bp for RNA-seq and single reads of 50 bp for ChIP-seq.

### ChIP-seq analysis

Raw reads were demultiplexed into FASTQ format allowing up to one mismatch using Illumina bcl2fastq2 v2.18. Reads were aligned to the human genome (hg38) using Bowtie2 (version 2.3.4.1) with default parameters (Langmead and Salzberg 2012). For samples with fly spike-in, reads were first mapped to the Drosophila melanogaster genome (dm6), and unmapped reads were then aligned to the human genome (hg38). Reads per million (RPM) normalized bigWig tracks were generated by extending reads to 150bp. For spike-in ChIP-seq data, we also generated spike-in normalized bigWig tracks (RPM normalization factor = 1E6 / number of reads aligned to hg38, and spike-in normalization factor = 1E6 / number of reads aligned to dm6).

### Peak calling

epic2 (with options: -gn hg38 -fs 200 -fdr 0.05) was used to call wide peaks for H3K36me3 ChIP-seq data for FL, N3, and N3ΔSRI (Stovner et al. 2019). Next, ChIPseeker was applied (with options: genomicAnnotationPriority=c(‘Intergenic’, ‘5UTR’, ‘3UTR’, ‘Exon’, ‘Intron’)) to obtain the genomic feature distribution (Ensembl 96 release) under peaks (Yu et al. 2015).

### Metagene Plots

14533 Protein-coding genes (Ensembl 96 release) were selected with length ≥ 600bp and no other genes within −2Kb TSS and +2Kb TES regions. Metagene regions were from −2Kb TSS to +2Kb TES. In addition, 2Kb upstream TSS and downstream TES regions are grouped into 100 bins (20bp per bin), respectively. The gene body region is grouped into 300 bins (at least 2bp per bin since the minimum gene length is 600bp). In total, each gene is grouped into 500 bins. The average normalized (RPM or spike-in) H3K36me3 signals in each bin were plotted using R package EnrichedHeatmap (Gu et al. 2018).

### H3K36me3 on exons and introns

Protein-coding genes were selected, and the longest transcript for each gene was chosen. Also, we removed any overlapping transcripts (ignore strand). As a result, 15311 transcripts were used to calculate the H3K36me3 signal (RPM normalized) distribution on exons/introns. The average exon/intron signal is defined as the total H3K36me3 signals on all exons/introns divided by the total exon/intron length.

### RNA-seq analysis

Raw reads were demultiplexed into FASTQ format allowing up to one mismatch using Illumina bcl2fastq2 v2.18. Reads were aligned to the human genome (hg38 and Ensembl 96 gene models) using STAR (version STAR_2.6.1c) (Dobin et al. 2013). TPM expression values were generated using RSEM (version v1.3.0) [5]. edgeR (version 3.24.3 with R 3.5.2) was applied to perform differential expression analysis, using only protein-coding and lncRNA genes (Robinson et al. 2009). To perform differential splicing analysis, we used rMATs (version 4.0.2) with default parameters starting from FASTQ files (Shen et al. 2014). FDR cutoff of 0.05 was used to determine statistical significance.

## ACCESSION NUMBERS

The data sets are available in the Gene Expression Omnibus (GEO) database under the accession number GSE147752.

## ACKNOWLEDGMENT

The authors are grateful to Dr. Kimryn Rathmell, Vanderbilt Institute for Infection, Immunology and Inflammation from providing SETD2 knock out 293T cells. The authors would like to thank the members of the Workman lab for their critical suggestions to improve the manuscript.

## AUTHOR CONTRIBUTION

S.B. conceptualized the work, designed and performed the experiments. S.B. wrote the manuscript. N.Z. and H.L. analyzed the high-throughput sequencing data and made the related plots. J.L.W conceived the idea of the work, provided supervision, acquired funding, and revised the manuscript.

