## supplementary information for "The disordered regions of SETD2 govern H3K36me3 deposition by regulating its proteasome-mediated decay"

(a)

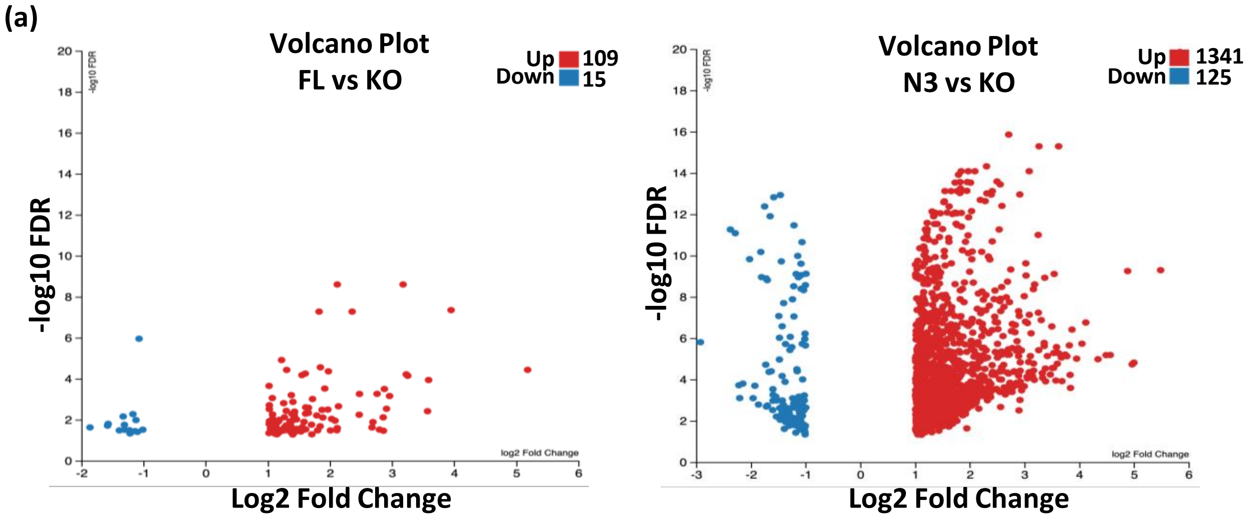


(b)

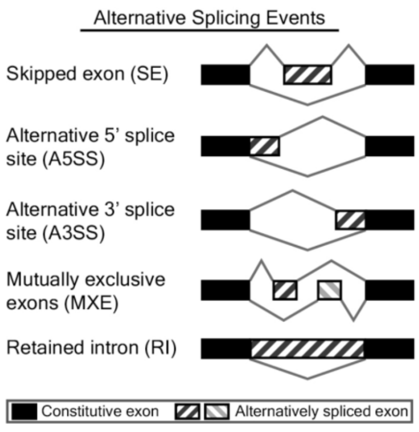


| Alternative Splicing Events | FL vs KO Up | FL vs KO Down | N3 vs KO Up | N3 vs KO Down |
|---|---|---|---|---|
| Skipped exon (SE) | 140 | 222 | 574 | 434 |
| Alternative 5' splice site (A5SS) | 6 | 18 | 104 | 51 |
| Alternative 3' splice site (A3SS) | 7 | 18 | 93 | 47 |
| Mutually exclusive exons (MXE) | 11 | 11 | 49 | 51 |
| Retained intron (RI) | 10 | 52 | 451 | 91 |
| | 174 | 321 | 1271 | 674 |

**Supplementary information S1:** (a) Volcano plot showing the expression changes occurring on SETD2 N3 and SETD2 FL expression in KO cells. Each dot is a gene. RNA was isolated 72 hours post transfection. (b) RMATS plot depicting the differential alternative splicing events in SETD2 N3 and SETD2 FL rescued KO cells. The cartoon was taken from http://rnaseq-mats.sourceforge.net/.

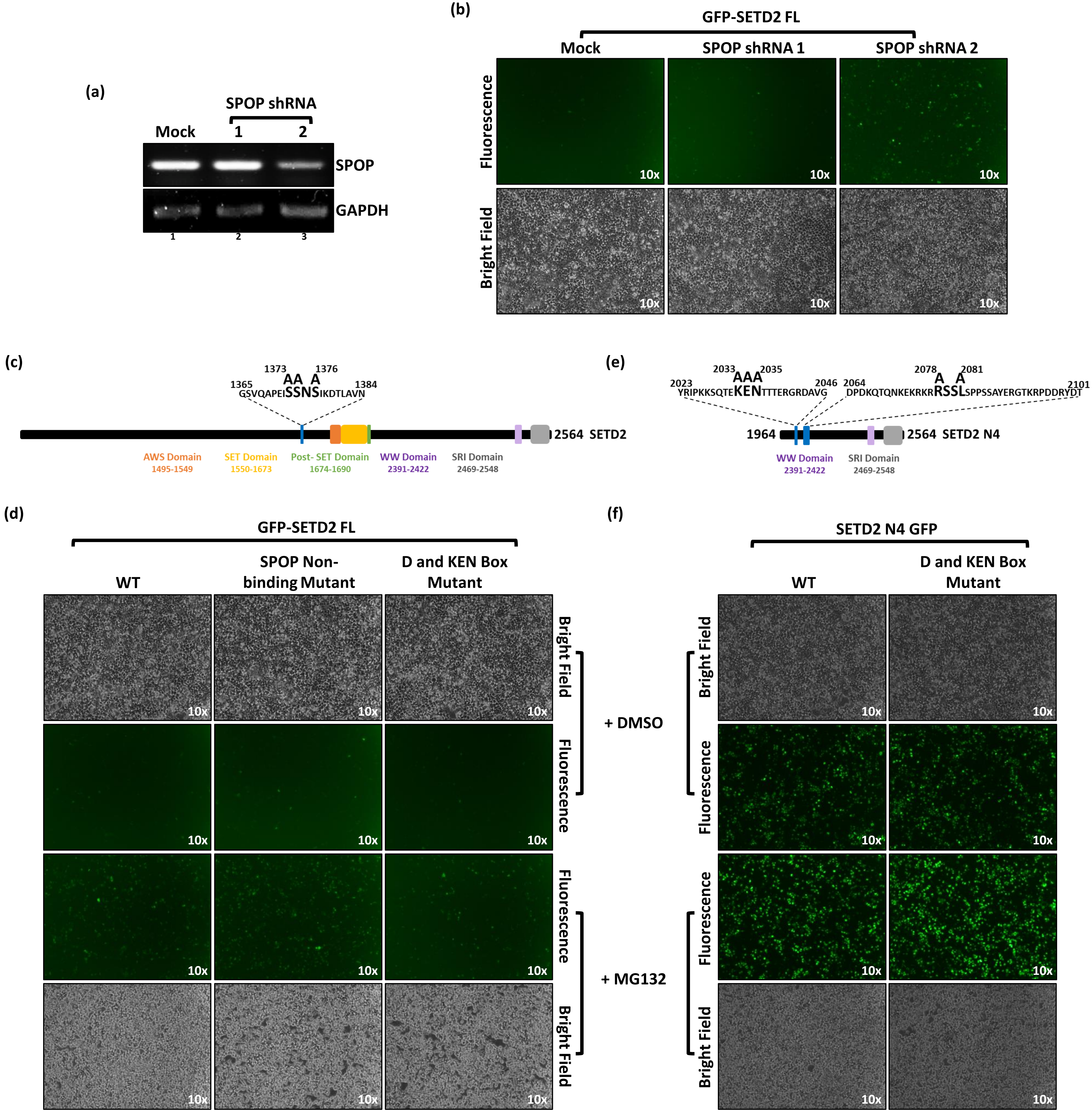
(a)
SPOP shRNA
Mock
1
2
SPOP
GAPDH
(b)
GFP-SETD2 FL
Mock
SPOP shRNA 1
SPOP shRNA 2
Fluorescence
Bright Field
10x
(c)
1365
1373
1376
1384
AA A
GSVQAPEISSNSIKDTLAVN
2564 SETD2
AWS Domain
1495-1549
SET Domain
1550-1673
Post- SET Domain
1674-1690
WW Domain
2391-2422
SRI Domain
2469-2548
(e)
2023
2033
2035
2046
2064
2078
2081
2101
AAA
A A
YRIPKKSQTEKENTTTERGRDAVG
DPDKQTQNKEKRKRRSSLSPPSSAYERGTKRPDDRYDT
1964
2564 SETD2 N4
WW Domain
2391-2422
SRI Domain
2469-2548
(d)
GFP-SETD2 FL
WT
SPOP Non-binding Mutant
D and KEN Box Mutant
Bright Field
Fluorescence
+ DMSO
+ MG132
(f)
SETD2 N4 GFP
WT
D and KEN Box Mutant

**Supplementary information S2:** (a) RNA was isolated from GFP-SETD2 FL expressing 293T cells transfected with SPOP shRNA and RT-PCR was performed to check transcript levels. GAPDH was used as normalization control. (b) Microscopy images showing effect of SPOP shRNA treatment on expression of GFP-SETD2 FL in 293T cells. (c, e) Cartoon depicting the position and sequence of SPOP binding and putative D/KEN Box in SETD2 and the mutation introduced to disrupt those. (d, f) Microscopy images showing expression of GFP-SETD2 constructs in 293T cells and the effect of MG132 treatment on their expression. WT- Wild Type.

#### Supplementary information S2 Results

SPOP was knocked down by shRNA in GFP-SETD2 FL-expressing 293T cells. Of the two shRNAs used, shRNA #2 treatment reduced SPOP transcript levels [supplementary information S2a]. Depletion of SPOP led to only a modest increase in GFP-SETD2 FL accumulation in 293T cells [supplementary information S2b]. To further substantiate this result, an SPOP Non-binding Mutant M3 of GFP-SETD2 FL was made [supplementary information S2c]. A significant increase was not observed in the expression level of the mutant compared to the WT [supplementary information S2d].

Next, we looked at the possible role of APC in regulating SETD2 degradation. The APC recognizes the D and KEN Box in a protein and targets them for degradation by the UPS. SETD2 has a putative D Box at position 2033-2035 and KEN Box at 2078-2081 [supplementary information S2e]. Both these regions were mutated in GFP-SETD2 FL and the SETD2 fragment GFP-SETD2 N4 (1964-2564), where the D and KEN boxes are located. The expression of these mutants was tested in 293T cells [supplementary information S2d, e and f]. Comparison of the mutant with the WT expression did not reveal any discernible difference. In addition, both the GFP-SETD2 FL SPOP Non-binding Mutant M3 and the D-KEN box mutants retained sensitivity to MG132 [supplementary information S2d and f].

(a)

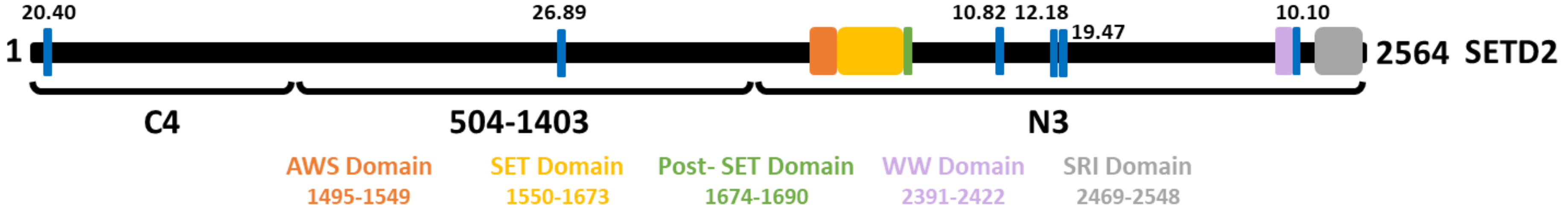


http://emboss.bioinformatics.nl/cgi-bin/emboss/epestfind

(b)

| Position | Sequence | Score |
|---|---|---|
| 20-31 | HPTPEEEENEAK | 20.40 |
| 96-115 | KQSDTPNPPAVPLQVDSTPK | 5.17 |
| 117-134 | KMEIGDTLSTAEESSPPK | 8.87 |
| 169-216 | HAAPLPAVIAESTTVDSPPSSPPPPPPPAQ ATTLSSPAPVTEPVALPH | 9.62 |
| 1035-1063 | HEDYSGSSESSNDESDSEDTDSDDSSIPR | 26.89 |
| 1074-1089 | KNSTLPMEETSPCSSR | 6.78 |
| 1831-1850 | KTAVPPLSEGDGYSSENTSR | 5.5 |
| 1852-1863 | HTPLNTPDPSTK | 8.28 |
| 1863-1874 | KLSTEADTDTPK | 10.82 |
| 1948-1963 | KLPTSEPEADAEIEPK | 12.18 |
| 1969-1994 | KLEEPINEETPSQDEEEGVSDVESER | 19.47 |
| 2413-2433 | RQTQWDPPTWESPGDDASLEH | 10.12 |

(c)

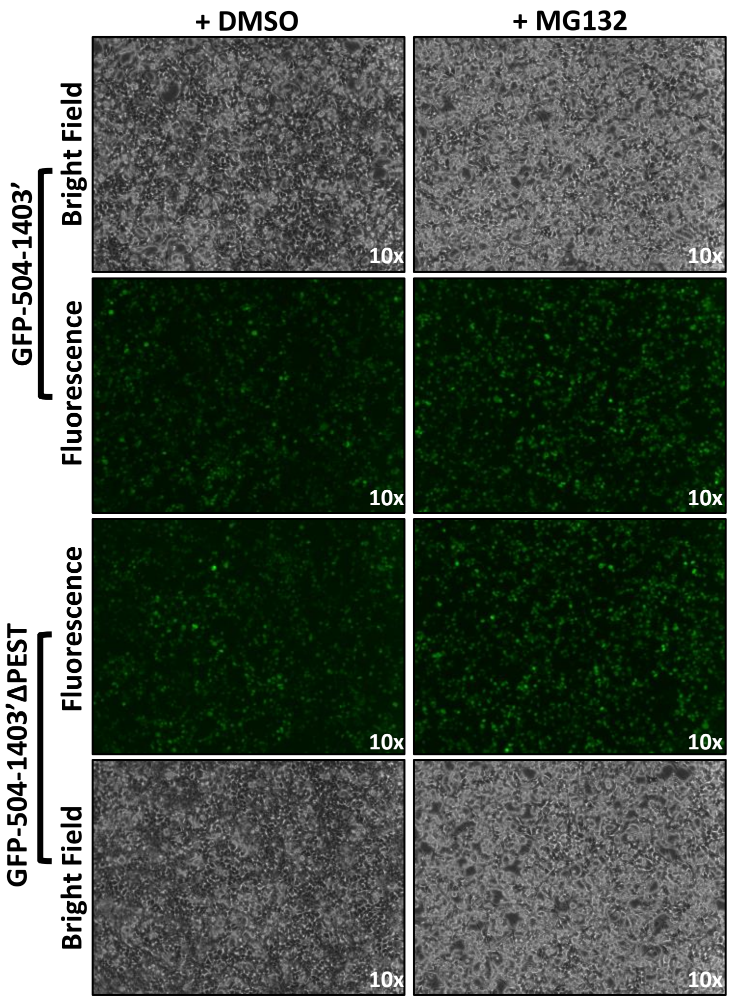


(d)

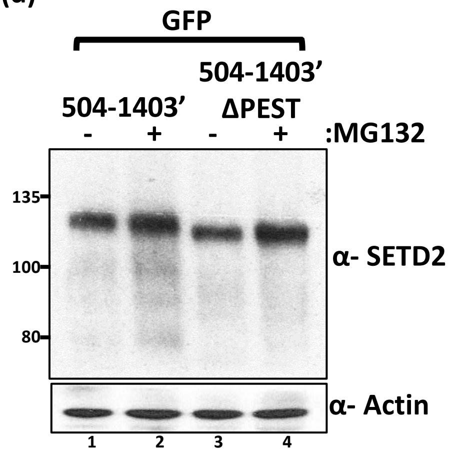


**Supplementary information S3:** (a) Cartoon illustrating the location of PEST motifs in SETD2 of score >10 (>5 is considered significant). (b) Position, sequence and score of PEST motifs in SETD2. (c) Microscopy images showing expression of GFP-SETD2 fragments. See text for more details. (d) Western blot of whole-cell lysates of cells shown in (c) probed with the depicted antibodies.

### Supplementary information S3 Results

Sequence analysis of SETD2 revealed the presence of multiple PEST motifs across its sequence with a significant score of more than 5 [supplementary information S3a, b]. The highest score of 26.89 was found in the stretch of 504-1403 residues [supplementary information S3a]. To test the possible contribution of PEST motifs in governing SETD2 degradation, PEST motif from C-Myc NLS containing GFP-504-1403 (504-1403’) was deleted and its expression was checked in 293T cells by fluorescence microscopy and western blotting of whole cell extracts with anti-SETD2 antibody [supplementary information S3c, d]. The expression of the segment GFP-504-1403’ with and without the PEST motif were similar, suggesting that the SETD2 IDRs have a direct effect on its half-life rather than embedding a destruction signal. This is consistent with a study that found that disordered segments have direct effects on half-life by forming initiation sites for degradation by the proteasome rather than acting indirectly by embedding destruction signals. However, the possibility that other PEST motifs may govern SETD2 stability cannot be ruled out from our experiments.

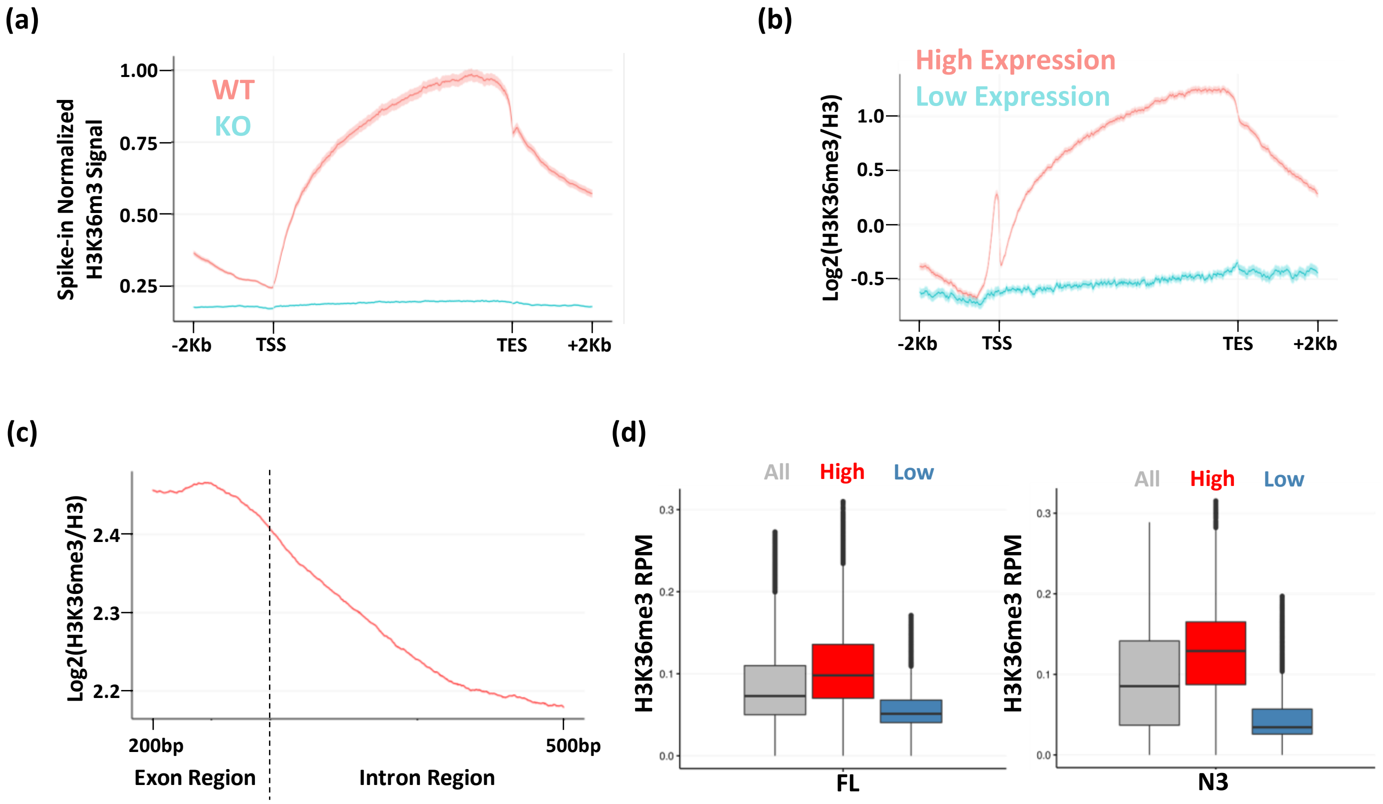


**Supplementary information S4:** (a) Metagene plot depicting the spike-in normalized distribution of H3K36me3 in WT and KO (setd2Δ) 293T cells. (b) Metagene plot showing the H3 normalized distribution of H3K36me3 over high/low expressed genes. High expression is TPM >= 4 and low expression if 0 < TPM <= 2 in WT samples. (c) Metagene plot showing the H3 normalized distribution of H3K36me3 over exon and introns of expressed genes. (d) Box plot showing the H3K36me3 and expression level of genes. The expression cutoffs are 0 < TPM <= 2 and TPM >= 4 to define low- and high-expressed protein coding genes, respectively, in FL and N3 samples; and 0 < TPM <= 0.1 and TPM >= 0.5 to define low- and high-expressed lncRNA genes, respectively, in FL and N3 samples.

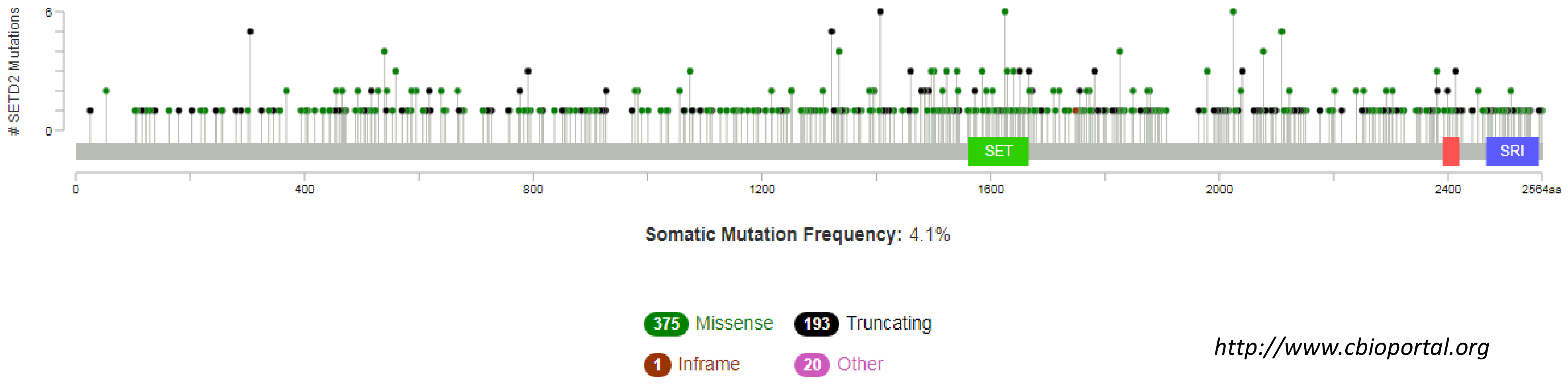


**Supplementary information S5:** Cartoon depicting the location of various mutations found in SETD2 gene of cancer patients that may result in the generation of truncated forms of SETD2 with altered half-lives.

| Oligo | Sequence (5’-3’) |
|---|---|
| SPOP_F | TACCCTCTTCTGCGAGGTGA |
| SPOP_R | CGGGAATTCTCCCACAGTCC |
| GAPDH _F | TTCGACAGTCAGCCGCATCTTCTT |
| GAPDH _R | CAGGCGCCCAATACGACCAAATC |
| SPOP_shRNA_1 F | CCGGGATTCAAGAAATTCATCCGTAGATATCTACGGATGAATTTCTTGAATCTTTTTG |
| SPOP_shRNA_1 R | AATTCAAAAAGATTCAAGAAATTCATCCGTAGATATCTACGGATGAATTTCTTGAATC |
| SPOP_shRNA_2 F | CCGGTTCCAGGCTCACAAGGCTATCGATATCGATAGCCTTGTGAGCCTGGAATTTTTG |
| SPOP_shRNA_2 R | AATTCAAAAATTCCAGGCTCACAAGGCTATCGATATCGATAGCCTTGTGAGCCTGGAA |

**Supplementary information S6:** Sequence of oligos used to perform RT-PCR and knockdown.
